# Structural characterization of Nonstructural protein 1 from SARS-CoV-2

**DOI:** 10.1101/2020.09.08.288191

**Authors:** Cameron Semper, Nobuhiko Watanabe, Alexei Savchenko

## Abstract

Severe acute respiratory syndrome (SARS) coronavirus-2 (SARS-CoV-2) is a single-stranded, enveloped RNA virus and the etiological agent of the current COVID-19 pandemic. Efficient replication of the virus relies on the activity of nonstructural protein 1 (Nsp1), a major virulence factor shown to facilitate suppression of host gene expression through promotion of host mRNA degradation and interaction with the 40S ribosomal subunit. Here, we report the crystal structure of the globular domain of SARS-CoV-2 Nsp1, encompassing residues 13 to 127, at a resolution of 1.65 Å. Our structure features a six-stranded, capped β-barrel motif similar to Nsp1from SARS-CoV and reveals how variations in amino acid sequence manifest as distinct structural features. Through comparative analysis of structural homologues, we identified a topological signature associated with this protein fold that facilitated modeling of Nsp1 from MERS-CoV. Combining our high-resolution crystal structure with existing data on the C-terminus of Nsp1 from SARS-CoV-2, we propose a model of the full-length protein. Our results provide unparalleled insight into the molecular structure of a major pathogenic determinant of SARS-CoV-2.

## Introduction

In March of 2020, the World Health Organization (WHO) declared Coronavirus Disease 2019 (COVID-19) a global pandemic. As of September 2020, there have been more than 26,000,00 cases of infection reported globally and approximately 860,000 deaths attributed to COVID-19 [1]. The etiological agent of this pandemic has been identified as Severe Acute Respiratory Syndrome coronavirus-2 (SARS-CoV-2), a member of the *Betacoronavirus* genus and closely related to the SARS-CoV that caused the SARS outbreak of 2002-2004 [2, 3].

Coronaviruses infect a diverse array of vertebrates, with infection typically resulting in respiratory disease or gastroenteritis [4]. Their broad host range allows for a substantial reservoir for human infection and mutation facilitates cross-species transmission [5]. There is mounting evidence that SARS-CoV-2 originated via mutation and cross-species transmission of a pangolin coronavirus with which it shares ~97% sequence identity [6]. This highlights the dramatic impact that relatively few amino acid substitutions can have in coronaviruses and underscores the urgent need for characterization of these infectious agents at a molecular level.

SARS-CoV-2 is an enveloped, positive-sense RNA virus with a single-stranded genome approximately 30 kb in size. The genome is 5’-capped and 3’poly-adenylated and the first ~2/3 of the genome encode two overlapping reading frames that produce polyprotein 1a and 1ab [7]. Downstream of this region, the remaining 1/3 of the genome encodes for structural proteins and a number of ORFs that produce accessory proteins largely of unknown function. Polyprotein 1a and 1ab are large polypeptides that are processed post-translationally by viral-encoded proteases to produce non-structural proteins (nsp) 1-16 [8]. The genome also contains 5’ and 3’ untranslated regions (UTRs), the former of which plays a critical role in self-recognition that allows for SARS-CoV-2 protein production to occur unabated while host gene expression is suppressed [9]. At the core of this mechanism is Nsp1, a 180 amino acid (AA) protein produced via processing of polypeptide 1a and 1ab by the Papain-like protease domain of Nsp3 (Figure 1A) [10].

**Figure 1.**
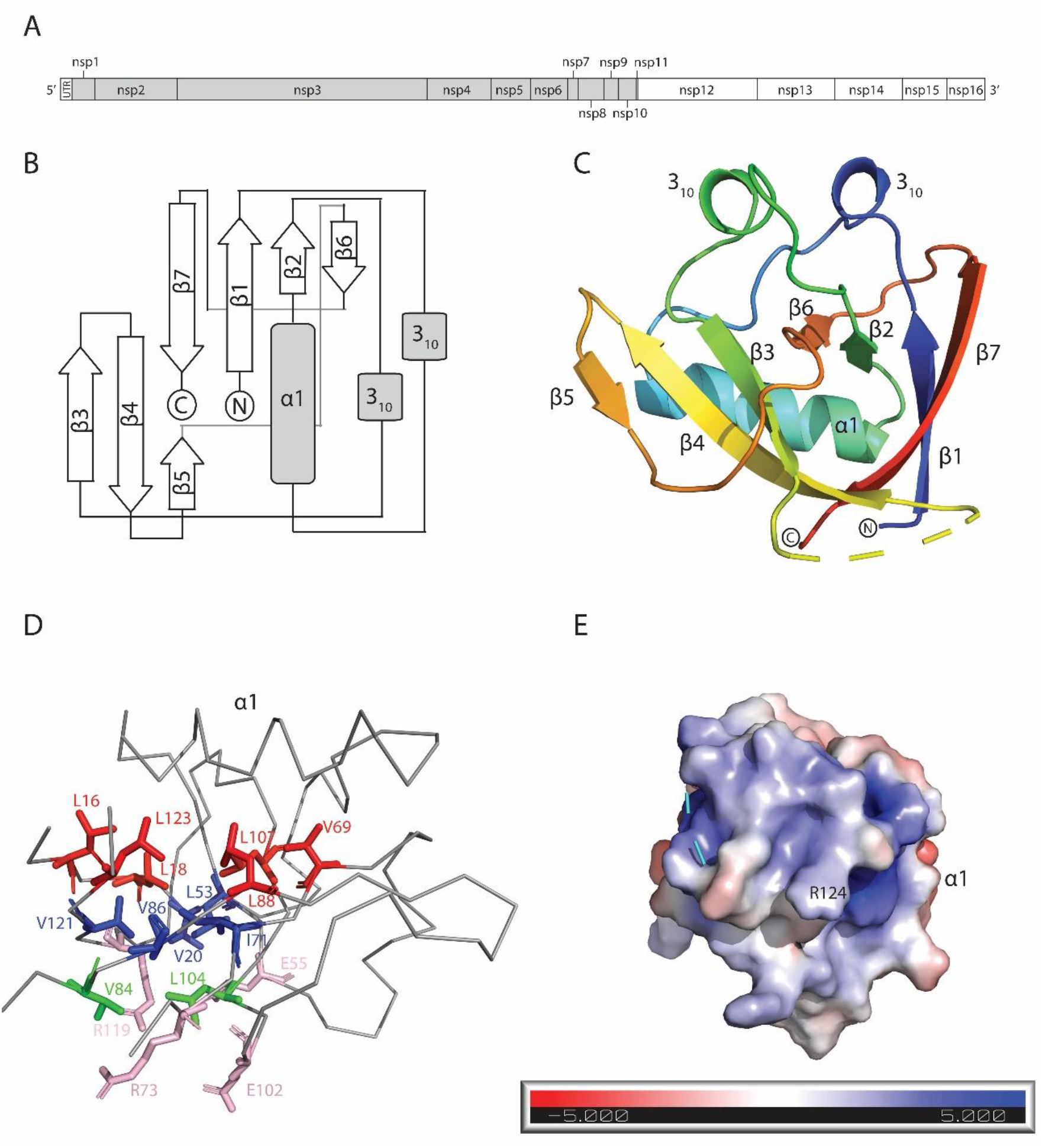
The structure of SARS-CoV-2 Nsp1_13-127_. (A) Schematic of Orf1ab from SARS-CoV-2. The grey shading encompasses the non-structural proteins produced by Orf1a. (B) Topological arrangement of SARS-CoV-2 Nsp1_13-127_. (C) Cartoon representation of the SARS-CoV-2 Nsp1_13-127_ structure coloured from N-terminus (blue) to C-terminus (red). Secondary structure elements and the N- and C-termini are labeled. (D) Ribbon depiction of the structure, with sidechains that contribute to the hydrophobic core of the beta barrel shown in stick representation. The three layers of side chains, plus the charged residues at the bottom are coloured and the residues within the layers are labeled. (E) Electrostatic surface potential of the likely RNA-binding interface of SARS-CoV-2 Nsp1_13-127_. R124 which is critical for the interaction with SL1 of the 5’UTR is labeled.

Nsp1 mediates a two-pronged approach to suppression of host gene expression. Firstly, it inhibits translation of host proteins during the initiation stage through interaction with the 40S ribosomal subunit. Secondly, Nsp1 promotes the degradation of host mRNA by endonucleolytic cleavage within the 5’UTR, which in turn leads to accelerated Xrn1-mediated mRNA decay [11, 12]. Viral mRNAs are able to avoid the fate of host mRNAs through interaction between Nsp1 and the stem-loop 1 (SL1) motif found in the viral 5’ UTR [9]. Nsp1 from SARS-CoV has been shown to inhibit the Type I Interferon response in infected cells, allowing for the virus to circumvent the innate immune response [13]. Expression of SARS-CoV Nsp1 has also been shown to induce the production of chemokines, suggesting this protein may play a role in the “cytokine storm”, a maladaptive release of cytokines in response to infection, associated with a number of COVID-19 infections [14, 15]. Thus, Nsp1 has emerged as a major pathogenicity factor that plays a critical role in the coronavirus infection cycle [16]. Deletion or mutation of nsp1 results in viral attenuation in infection models and restores the innate immune response in infected cells [17]. Based on its central role in suppression of the host immune response and essentiality to infection, Nsp1 has been proposed as a therapeutic target for the treatment of COVID-19 [16]. A prerequisite to any investigation into possible interventions targeting Nsp1 is high-resolution structural data that can facilitate robust *in silico* screening. A partial structure corresponding to the N-terminal fragment of Nsp1 from SARS-CoV was resolved by NMR spectroscopy and revealed a unique β-barrel motif. The corresponding domain from SARS-CoV-2 shares 86% sequence identity with its SARS-CoV ortholog, which is a considerable level of sequence diversity compared to many other non-structural proteins encoded by the SARS-CoV-2 genome (e.g. Nsp12 shares 96% identity with SARS-CoV ortholog [18]). This highlights the need for a pursuit of structural characterization of SARS-CoV-2 proteins.

Here, we report the high-resolution crystal structure of the globular N-terminal domain of Nsp1 from SARS-CoV-2 at 1.65 Å resolution. Our data reveals a high level of structural conservation between Nsp1 of SARS-CoV-2 and SARS-CoV, but also some unique structural features that likely contribute to increased stability of the β-barrel fold in SARS-CoV-2 Nsp1. Comparative analysis reveals additional structural homologues in Nsp1 proteins from *Alphacoronaviruses*, despite low levels of shared sequence identity. These results highlight the critical role this unique protein fold plays in facilitating viral infection and suppression of host gene expression.

## Results

### Crystal structure of SARS-CoV-2 Nsp1_13-127_

In pursuit of structural characterization of Nsp1 from SARS-CoV-2 we expressed a codon optimised version of the ORF encoding this protein in *E. coli*. Using this expression system, we obtained purified full-length Nsp1 after a two-step purification protocol (see details in Material and Methods). However, we were unable to crystallize the full-length Nsp1 protein, likely due to the presence of flexibly disordered regions at the N- and C-termini.

Previous studies of Nsp1 from SARS-CoV reported the presence of a distinct globular domain comprised of residues 13 to 127 [19]. To structurally characterise the N-terminal fragment of Nsp1 from SARS-CoV-2 (Nsp1_13-127_), we sub-cloned it and expressed it in *E. coli*. The expression level of Nsp1_13-127_ was comparable to that of the full-length protein and with this fragment we obtained well-diffracting crystals in several conditions that enabled structural determination of this portion of Nsp1.

The crystal structure of SARS-CoV-2 Nsp1_13-127_ was determined via molecular replacement using the SARS-CoV Nsp1(PDB: 2HSX) solution structure as the search model and refined to a resolution of 1.65 Å (Table 1). The asymmetric unit contained a single molecule of Nsp1_13-127_. Analysis of the Nsp1_13-127_ structure by PDBePISA revealed the protein has a total solvent-exposed area of 6102 Å^2^ [20].

**Table 1.** X-ray crystallographic statistics.

|  | SARS-Cov-2<br>Nsp1<sub>13-127</sub> |
| --- | --- |
| **Data collection** |  |
| Space group | *P*4<sub>3</sub>2<sub>1</sub>2 |
| Cell dimensions |  |
| *a*, *b*, *c* (Å) | 30.0, 37.0, 144.8 |
| α, β, γ (°) | 90.0, 90.0, 90.0 |
| Wavelength | 0.97913 |
| Resolution (Å) | 50.0-1.65<br>(1.68-1.65) |
| *R*<sub>meas</sub><sup>1</sup> | 0.058 (1.091) |
| *R*<sub>pim</sub><sup>2</sup> | 0.020 (0.387) |
| *I* / *sI* | 35.6 (2.0) |
| Completeness (%) | 99.8 (99.5) |
| Redundancy | 8.2 (7.4) |
| CC1/2 | 0.997 (0.690) |
| **Refinement** |  |
| Resolution (Å) | 32.95-1.65 |
| No. reflections | 12884 |
| *R*<sub>work</sub> / *R*<sub>free</sub><sup>3</sup> | 0.203/0.248 |
| No. atoms |  |
| Protein | 828 |
| Water | 115 |
| Average *B*-factors (Å<sup>2</sup>) | 20.7 |
| Protein | 19.2 |
| Water | 31.2 |
| R.m.s deviations |  |
| Bond lengths (Å) | 0.006 |
| Bond angles (°) | 0.872 |
| Ramachandran (%) |  |
| Favoured | 97.1 |
| Allowed | 2.9 |
| Disallowed | 0 |

\*Values in brackets refer to highest resolution shells.
1. $R_{\text{meas}} = \frac{\sum_{hkl} \sqrt{(n/n-1) \sum_{j=1}^n |I_{hkl,j} - \langle I_{hkl} \rangle| / \sum_{hkl} \sum_j I_{hkl,j}}}{\dots}$
2. $R_{\text{pim}} = \frac{\sum_{hkl} \sqrt{(1/n-1) \sum_{j=1}^n |I_{hkl,j} - \langle I_{hkl} \rangle| / \sum_{hkl} \sum_j I_{hkl,j}}}{\dots}$
3. $R = \frac{\sum |F_p^{\text{obs}} - F_p^{\text{calc}}| / \sum F_p^{\text{obs}}}{\dots}$ , where $F_p^{\text{obs}}$ and $F_p^{\text{calc}}$ are the observed and calculated structure factor amplitudes, respectively.

As in the case of SARS-CoV, the structure of SARS-CoV-2 Nsp1_13-127_ features a unique topological arrangement resulting in the formation of a six stranded (n = 6) β-barrel that is primarily antiparallel, with the exception of strands β1 (Q15 – V20) and β2 (C51 – V54) (Figure 1B). Additional major structural features include α1 (V35 – D48) helix, which is positioned as a cap along one opening of the β-barrel, two 310 helices that run parallel to each other and the β5 strand (I95 – Y97) which is not part of the β-barrel but forms a β-sheet interaction with the β4 strand (V84 – L92) (Figure 1C).

The core of the β-barrel is highly hydrophobic and is mainly comprised of the side chains of thirteen amino acids organized into three layers. The first layer, which is adjacent to α1 helix, is formed by sidechains of residues L16, L18, V69, L88, L107 and L123 (Figure 1D). The opening of the β-barrel at this layer is obstructed by the α1 helix, which contributes the side chain of L46 to the centre of this first layer of core hydrophobic residues. The middle layer of the β-barrel is comprised of the side chains of residues V20, L53, I71, V86 and V121, while the bottom layer features residues V84 and L104. Adjacent to this bottom layer are four solvent-exposed residues; E55, R73, E102 and R119 whose sidechains point inward towards the core of the β-barrel (Figure 1D). Consequently, the hydrophobic core at both ends of the β-barrel is occluded: by the α1 helix on one side and the charged residues (E54, R73, E102, R199) on the other.

A distinctive feature of Nsp1 that is evident in the crystal structure of Nsp1_13-127_ is the large number of flexible loops. For one of these loops (A76 – H81), we were unable to resolve the structure due to a lack of interpretable electron density. This region has been previously shown to be highly flexible as evidenced by the multitude of backbone confirmations observed in the NMR structure of SARS-CoV Nsp1 [19]. The positions of other loops in the crystal structure were stabilized via interactions between specific secondary structure elements. Specifically, the two 310 helices are aligned in the crystal structure forming H-bonds between R24 and Q63. This interaction appears to stabilize the position of two of the largest loops in the crystal structure, the β1-α1 (L21 – S34) loop and the β2- β3 (E54 – P67) loop.

Electrostatic surface potential analysis of the SARS-CoV-2 Nsp1_13-127_ structure revealed several regions of charged residue colocalization [21]. The most electronegative and electropositive patches appear adjacent on the Nsp1_13-127_ surface, separated by a small electroneutral region corresponding to the α1 helix. The electropositive patch continues around the surface of the protein, where the R124 residue is localized (Figure 1E). This residue has been shown to be essential for the interaction between Nsp1 and SL1 of the 5’ UTR, implicating this charged surface in RNA binding [9].

### Comparison with SARS-CoV Nsp1

The overall fold of SARS-CoV-2 Nsp1_13-127_ was highly similar to that of the corresponding SARS-CoV Nsp1 fragment in accordance with the significant primary sequence identity shared between the two proteins (Figure 2A). However, detailed comparative analysis revealed several notable differences between the two structures. The SARS-CoV-2 structure of Nsp1_13-127_ features two 3_10_ helices, compared to the single one found in SARS-CoV Nsp1. The primary sequence corresponding to the second 3_10_ helix is completely conserved between SARS-CoV-2 and SARS-CoV Nsp1 proteins, suggesting this secondary structure element may be transient in nature or influenced by differences elsewhere in SARS-CoV-2 Nsp1_13-127_. The two 3_10_ helices interact with each other through H-bonds in SARS-CoV-2 Nsp1_13-127_ but this interaction is notably absent in the NMR structure of Nsp1 from SARS-CoV [19].

**Figure 2.**
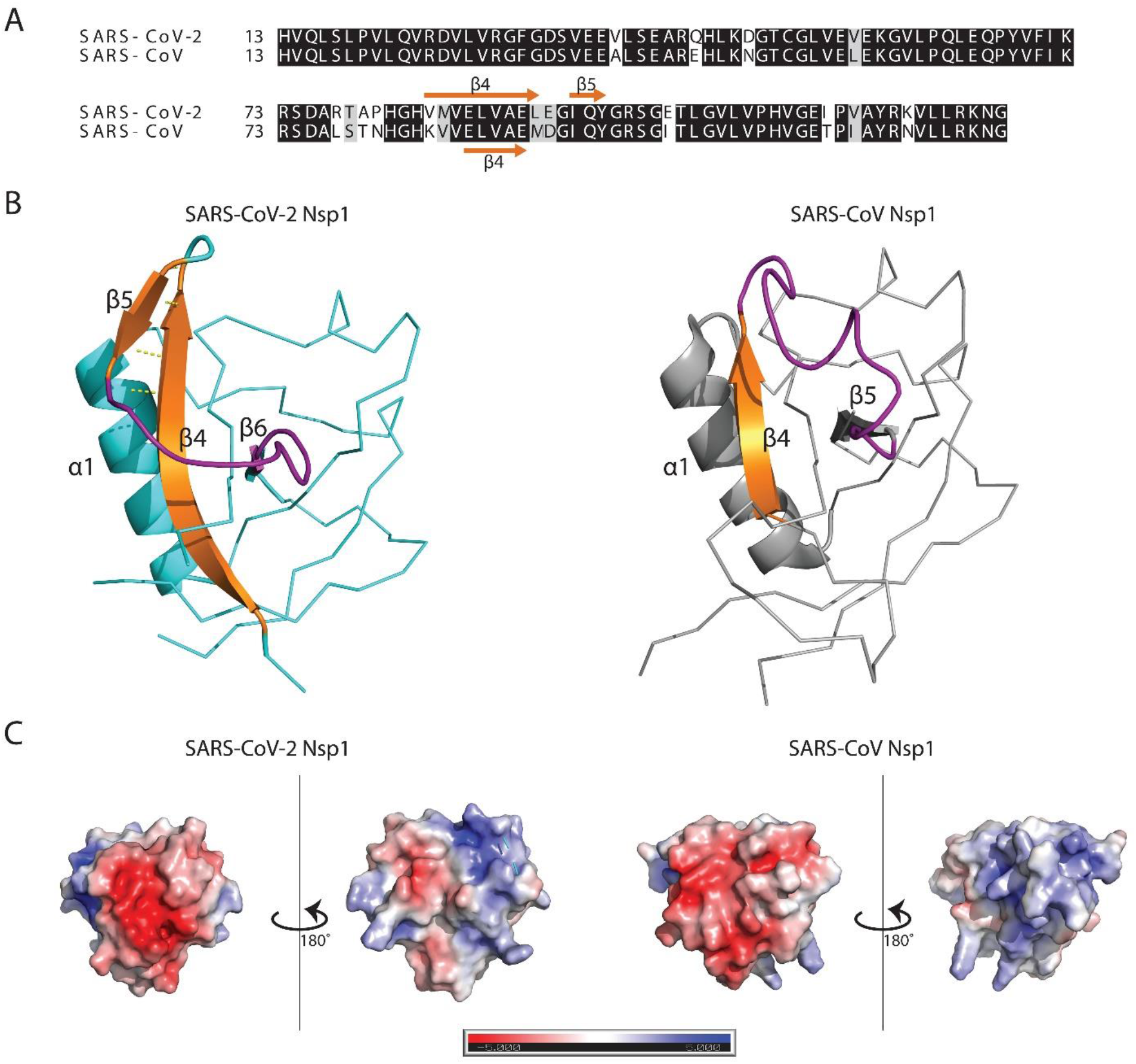
Comparison of SARS-CoV-2 Nsp1_13-127_ to SARS-CoV Nsp1. (A) Alignment of residues 13-127 of Nsp1 with conserved residues highlighted in black. (B) Ribbon representation of SARS-CoV-2 and SARS-CoV Nsp1_13-127_. β4 (shown in cartoon representation) is extended in the SARS-CoV-2 structure allowing for H-bonding with and formation of β5, which is absent in the SARS-CoV Nsp1structure. The loop region (and equivalent in SARS-CoV Nsp1) stabilized through interaction between β4 and β5 is coloured in purple. (C) Comparison of electrostatic surface potential of SARS-CoV-2 and SARS-CoV Nsp1_13-127_.

Two of the most striking differences between the two Nsp1 structures are the extension of β4 and the presence of an additional β-strand in the case of SARS-CoV-2 Nsp1_13-127_. Amino acid changes at positions 84 (K to V) and 85 (V to M) between SARS-CoV and SARS-CoV-2 Nsp1 result in an extension of β4 on the N-terminus by three residues (Figure 2B). This extension results in an increased number of H-bonds formed between β4 and β7 in SARS-CoV-2 Nsp1 compared to the pairing between β4 and β6 that occurs in SARS-CoV Nsp1. This extension also facilitates the formation of the β5 strand which is absent in SARS-CoV Nsp1. The β5 and β4 strands are part of a small β-sheet that stabilizes the loop region between β5 and β6 (β4 and β5 in SARS-CoV Nsp1) (Figure 2B).

Comparison of the electrostatic surface potential of Nsp1_13-127_ from SARS-CoV-2 and SARS-CoV shows additional differences between the two proteins. The SARS-CoV structure features a single patch of electronegative surface potential opposite a separate patch of electropositive surface potential. These two charged surfaces are oriented 180° opposite one another and there is a ring of electroneutral surface potential separating the two charged surfaces (Figure 2C). The SARS-CoV-2 Nsp1_13-127_ structure features an equivalent patch of electronegativity, but the electropositive surface is divided into two by a patch of mild electronegative surface potential.

Mapping the non-conserved residues onto the SARS-CoV-2 Nsp1_13-127_ structure revealed that most of the differences are in solvent-exposed residues. Distinct patches of variation are observed immediately adjacent to both ends of β4 and in the loop between β3 and β4 (S74 – H83), where 4 of 9 residues differ between SARS-CoV-2 and SARS-CoV (Figure 3A). The remaining sequence variation is broadly distributed across the Nsp1_13-127_ structure. Residues that contribute to the hydrophobic core of the β -barrel are conserved between SARS-CoV-2 and SARS-CoV, suggesting this particular organization is critical to the function of Nsp1.

**Figure 3.**
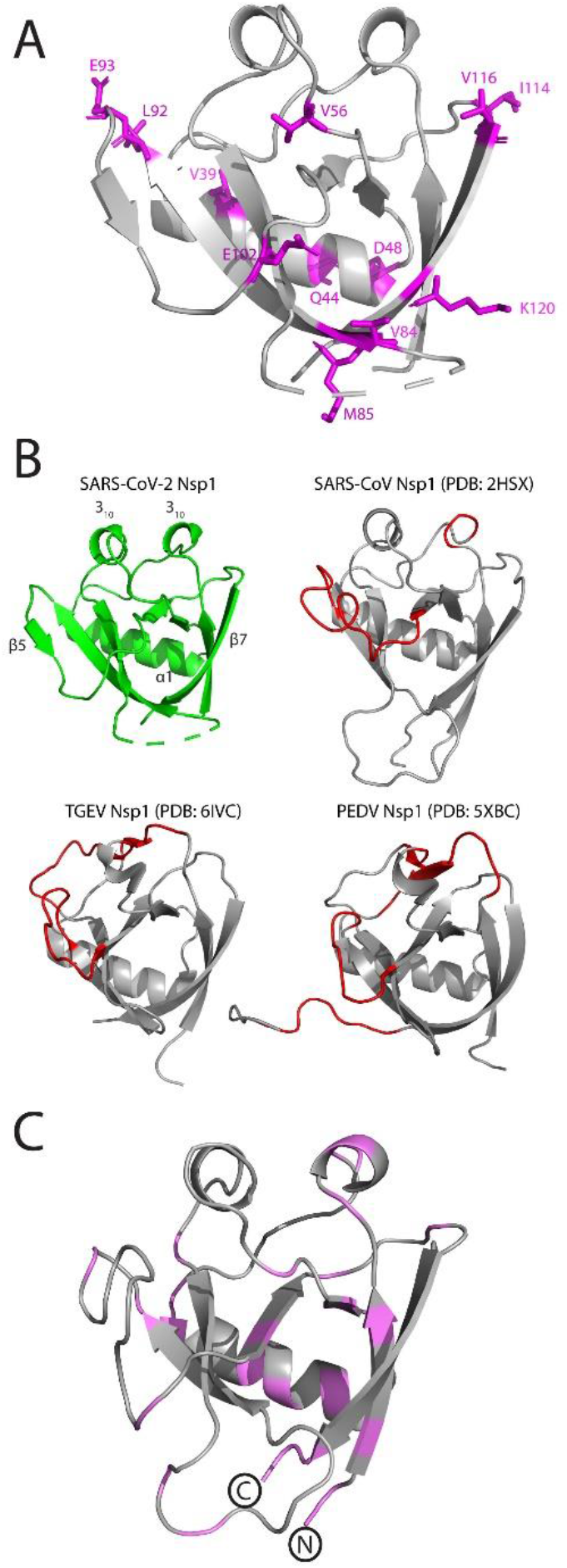
Comparison of the structure of SARS-CoV-2 Nsp1_13-127_ to structural homologues. (A) Cartoon representation of the SARS-CoV-2 Nsp1_13-127_ structure with non-conserved residues between SARS-CoV-2 and SARS-CoV highlighted in magenta. (B) SARS-CoV-2 Nsp1_13-127_ is shown in green, SARS-CoV Nsp1 (PDB: 2HSX), TGEV Nsp1 (PDB: 6IVC) and PEDV Nsp1 (PDB: 5XBC) are shown in grey with noticeable structural differences (compared to SARS-CoV-2) highlighted in red. Distinctive secondary structure features from the SARS-CoV-2 Nsp1_13-127_ structure are labeled. (C) Homology model of β-barrel domain of Nsp1 from MERS-CoV. Conserved residues between MERS-CoV and SARS-CoV-2 Nsp1 are highlighted in magenta and the N- and C-termini are labeled.

### Comparison with other structural homologues

Amino acid sequence-based searches for homologues of Nsp1 reveal a high level of sequence conservation among representatives of Lineage B of *Betacoronaviruses,* which includes several animal viruses closely related to SARS-CoV and SARS-CoV-2. Despite the presence of functionally equivalent Nsp1 proteins, no sequence-based homologues of SARS-CoV-2 Nsp1 were detected in other lineages of *Betacoronaviruses.* This lack of sequence conservation is observed even when comparing SARS-CoV-2 Nsp1 to other coronaviruses known to infect humans, including MERS, HCoV-OC43 and HCoV-HKU1.

A search for structural homologues using our high resolution structure of SARS-Cov-2 Nsp1_13-127_ identified three closely related structural homologues, all annotated as Nsp1 proteins (Table 2) [22]. Unsurprisingly, the Nsp1 structure from SARS-CoV was identified as the closest homologue, but the search also indicated strong similarity to the Nsp1 proteins from transmissible gastroenteritis virus (TGEV) and porcine epidemic diarrhea virus (PEDV). These *Alphacoronavirus* homologues superimposed with SARS-CoV-2 Nsp1_13-127_ with RMSD values of 2.8 Å and 2.9 Å over 114 and 118 Cα atoms, respectively (Table 2) [23]. Such appreciable levels of structural conservation, despite the low levels (<15%) of shared sequence identity, between these proteins highlights the plasticity of the unique protein fold found in Coronaviral Nsp1 proteins (Figure 3B).

**Table 2.** Structural homologues of SARS-Cov-2 Nsp1_13-127_.

| **PDB Code** | **DALI Z-Score (RMSD)** | **% Identity** | **Protein Name** | **Organism** |
| --- | --- | --- | --- | --- |
| 2GDT | 16.4 (1.6) | 88 | SARS-CoV<br>Nsp1 | Severe acute<br>respiratory syndrome<br>virus |
| 6IVC | 7.9 (2.8) | 14 | TGEV Nsp1 | Transmissible<br>gastroenteritis virus |
| 5XBC | 7.7 (2.9) | 15 | PEDV Nsp1 | Porcine epidemic<br>diarrhea virus |

To further examine the plasticity of this fold, we examined the predicted secondary structure of Nsp1 proteins from other coronaviruses known to infect humans (MERS, OC43, HKU1, 229E, NL63). These Nsp1 proteins share minimal primary sequence identity with Nsp1 from SARS-CoV-2 and a BLASTP search failed to identify them as homologues (Table S1). Despite the low primary sequence identity, we were able to identify topological fingerprints, regions comprised of 6-7 β-strands and at least one α-helix, in each of the Nsp1 sequences that are indicative of an arrangement capable of forming the capped β-barrel motif observed in the SARS-CoV-2 Nsp1_13-127_ crystal structure (Figure S1). We then modeled the region of MERS-CoV Nsp1 identified through this approach using our crystal structure as the threading template. This resulted in a high-scoring (C-score = 0.48) homology model featuring the capped β-barrel motif characteristic of the structure of SARS-CoV-2 Nsp1_13-127_ (Figure 3C). Mapping the modest primary sequence conservation between MERS-CoV and SARS-CoV-2 onto the homology model revealed it to be distributed throughout the structure (Figure 3C). The β-barrel in the MERS-CoV Nsp1 homology model retains the hydrophobic core observed in SARS-CoV-2, highlighting a level of functional conservation between the two proteins that may not have been predicted through sequence-based comparisons alone. At the C-terminus of the model is the R146/K147 dipeptide (R124/K125 in SARS-CoV-2) that has been shown to be involved in Nsp1-mediated RNA cleavage [24].

### Nsp1_13-127_ structure in context of the full-length SARS-CoV-2 Nsp1

Recent cryo-EM data of SARS-CoV-2 Nsp1 in complex with the 40S ribosome were able to resolve the structure of the C-terminal portion of Nsp1 at a resolution of 2.6 Å [25]. This study confirms the presence of a C-terminal α-helix spanning residues S166 to G179 and revealed a second helix spanning residues Y154 to N160 that was not predicted bioinformatically. The resulting helix-turn-helix motif interacts with the ribosome and anchors Nsp1 to the pre-initiation complex.

To place the crystal structure of Nsp1_13-127_ in context of full-length protein we analyzed the amino acid sequence using PSIPRED and DISPRRED3 [26, 27]. These tools predicted the N-terminal 12 residues outside of our structure contained a disordered protein-binding region and a largely unstructured C-terminal domain with the exception of a single predicted α-helix. The presence of large disordered regions adjacent to the globular domain of Nsp1 provide strong rationale for why the full-length protein failed to crystallize.

Using our crystal structure data in combination with insights into the C-terminal domain of Nsp1, we generated a homology-based model of the full-length Nsp1 protein. This model maintains the capped β-barrel motif observed in our crystal structure and features a largely disordered linker between the end of the globular domain and the C-terminal helix-turn-helix. The C-terminal domain is shown as extending outward from the core of the structure at the bottom of the β-barrel, at the face opposite of where the α1 helix localizes; however the disordered linker connecting this motif from the rest of the structure suggests that the position of this domain is likely to be highly dynamic and capable of multiple configurations (Figure 4). This predicted full-length structure aligns well with the partial density and model proposed by the recent cryo-EM structure of the SARS-CoV-2 Nsp1 C-terminus bound to the 40S ribosome [25].

**Figure 4.**
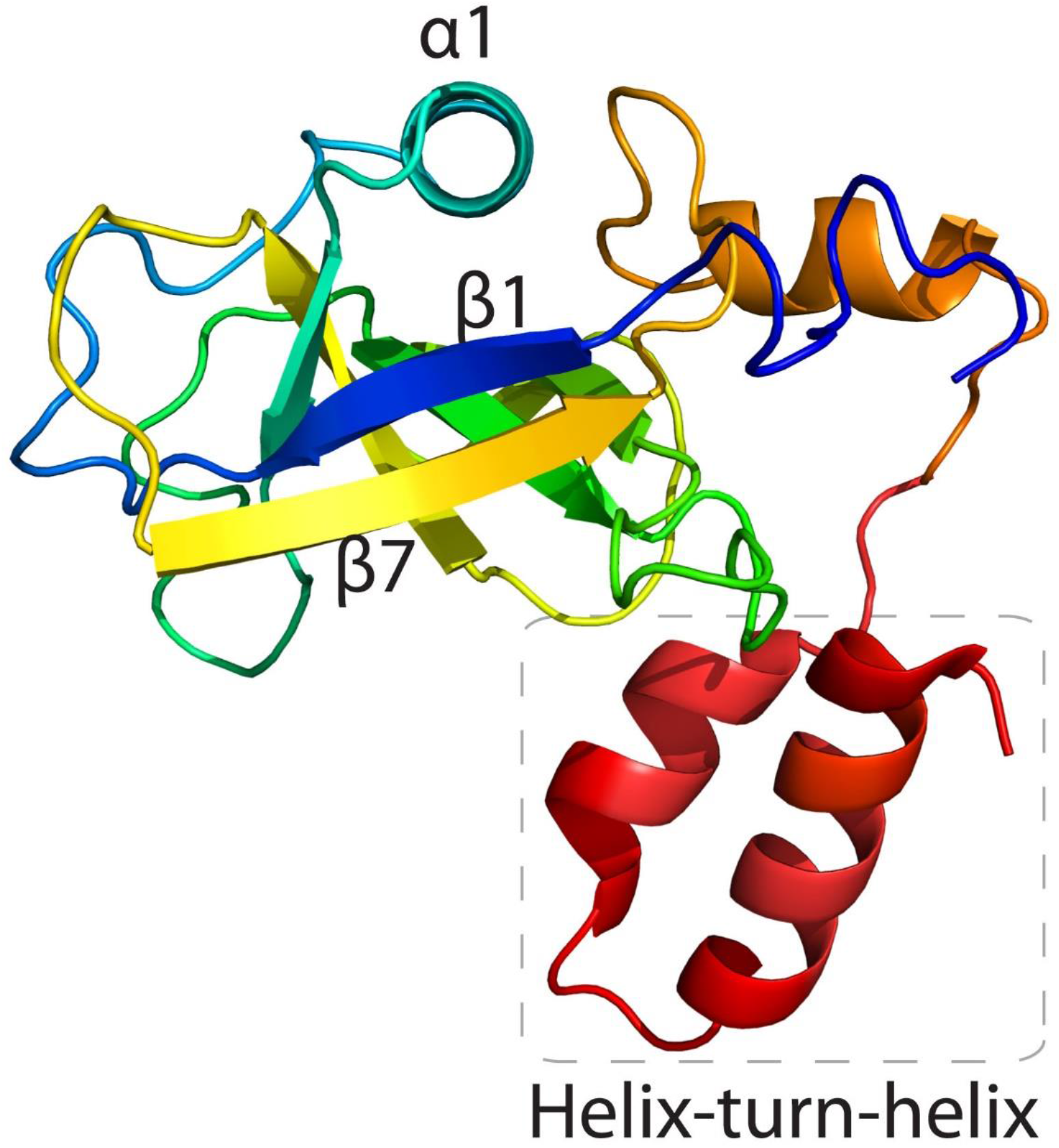
Model of full-length SARS-CoV-2 Nsp1. Cartoon representation of the model of the full-length SARS-CoV-2 Nsp1 coloured from N-terminus (blue) to C-terminus (red). Distinctive secondary structure features are labeled

## Discussion

SARS-CoV-2 Nsp1 arrests translation of host proteins through interaction with the 40S ribosome and in doing so suppresses the innate immune response to viral infection. Through interaction with SL1 in the viral 5’UTR, Nsp1 also distinguishes viral transcripts from host transcripts and allows for viral gene expression to continue unabated, highlighting the critical role this nonstructural protein plays in SARS-CoV-2 pathogenesis. Nsp1 proteins encoded by representatives of the *Coronaviridae* family feature significant amino acid sequence diversity, highlighted by the inability to identify homologues via BLAST of SARS-CoV-2 Nsp1 outside of the B lineage of *Betacoronaviruses.* Leveraging our high-resolution structural data, we were able to identify additional structural homologues of SARS-CoV-2 Nsp1, Nsp1 of TGEV and PEDV, with which it shares low levels of sequence identity. This result is in accordance with the recent data demonstrating that Coronaviral Nsp1 proteins exhibit similar function in suppression of host translation [24]. These observations suggest that the distinctive features of the SARS-CoV-2 Nsp1, highlighted by our structural data, may be critical to its function in self-recognition, via interaction with SL1, of viral RNAs and its role in promoting the degradation of host transcripts. This also provides insight into molecular features that could be expected in functionally similar proteins found in other important human pathogens, such as Nsp1 from MERS coronavirus.

Our sequence and structural based queries using SARS-CoV-2 Nsp1were only able to identify homologues within the *Coronaviridae* family. This raises the possibility that the specific fold and functions of Nsp1 are unique to Coronaviruses; however, the failure of sequence-based searches to identify Nsp1 from TGEV and PEDV as homologues of SARS-CoV-2 Nsp1 highlights the need for continued structural characterization of proteins involved in viral pathogenesis.

## Materials and Methods

### Cloning, expression and purification of SARS-CoV2 Nsp1

The portion of Orf1a encoding full-length Nsp1 was codon-optimized for *E. coli* expression, synthesized (Codex DNA) and cloned into the pMCSG53 expression vector at the SspI site via Gibson assembly. The fragment encompassing amino acid residues 13-127 of Nsp1 was PCR amplified and cloned into the same vector via ligation-independent cloning. Plasmids were transformed into the *E. coli* strain BL21(DE3)-Gold for protein expression. The same procedure was used for purification of full-length Nsp1 and Nsp1_13-127_. Cells were grown at 37°C and 200 rpm to an OD600 of 0.8, cooled to 20°C then induced with 1 mM IPTG and incubated for 16 hours. Cells were harvested via centrifugation at 5000 x *g*, resuspended in binding buffer (300 mM NaCl, 50 mM HEPES pH 7.5, 5 mM imidazole, 5% glycerol) and lysed via sonication. Lysates were centrifuged at 20,000 x *g* for 45 minutes at 4°C and the supernatant was incubated with Ni-NTA resin and rotated for 1 hour at 4°C. Nsp1 and Nsp1_13-127_ were eluted in elution buffer (300 mM NaCl, 50 mM HEPES pH 7.5, 250 mM imidazole, 5% glycerol) then incubated with Tobacco-etch virus (TEV) protease overnight to cleave the N-terminal polyhistidine tag while dialyzing to remove imidazole. The proteins were then passaged over a second Ni-NTA to remove impurities. Nsp1_13-127_ was immediately dialyzed into precrystallization buffer (300 mM NaCl, 10 mM HEPES pH 7.5), while full-length Nsp1was further purified via gel filtration using a Superdex75 column (300 mM NaCl, 10 mM HEPES pH 7.5).

### Crystallization

Crystals of Nsp1_13-127_ were grown at 298 K in 0.2 M sodium formate, 20% PEG3350 via the vapour diffusion sitting-drop method. Prior to data collection the crystals were soaked in 0.2 M sodium formate, 20% PEG3350, 30% glycerol and flash frozen in liquid nitrogen.

### Data collection, structure determination and refinement

X-ray diffraction data of crystals of Nsp1_13-127_ were collected at Advanced Photon Source Beamline 19 ID (Argonne, Illinoi USA) under cryo-stream at 93.15 K. Diffraction data were processed with HKL3000 suit [28]. Initial phase for Nsp1_13-127_ was obtained by molecular replacement with Molrep using NMR structure of SARS-CoV Nsp1 (PDB: 2HSX) as a search model [29]. Subsequently, the initial electron density map was improved though density modification by parrot and the model was built using buccaneer [29]. Final model was produced by the cycle of manual model building and refinement using Phenix.refine and Coot [30, 31]. All geometry was verified using Phenix validation tools (Ramachandran statistics: Favoured (97.1%), additionally allowed (2.9%), disallowed (0.0%)) and the wwPDB server.

### Structure analysis and homology-modeling

Electrostatic potential surfaces were calculated using Pymol using APBS[21]. · Structure similarity searches of the Protein Data Bank were performed using the Dali server [22].Secondary structure prediction of Nsp1 homologues was done using PSIPRED [26]. Homology modeling of MERS-CoV was performed by I-TASSER using the crystal structure of SARS-CoV-2 Nsp1_13-127_ as the threading template [32, 33]. The model produced by I-TASSER was subjected to further energy minimization using the YASARA energy minimization server to improve stereochemical property [34]. Stereochemical properties of the final model was validated with Ramachandran plot using PROCHECK server and 90% of residues are in the favoured/additionally allowed region [35]. The model of the full-length SARS-CoV-2 Nsp1 was generated using the Robetta structural predication server with the SARS-CoV-2 Nsp1_13-127_ structure defined as the template [36].

## Supporting information

Supplementary matherial for Semper at al.

## Acknowledgements

We thank Changsoo Chang and the Structural Biology Center Team at APS for data collection. The structure presented was solved as part of the Center for Structural Genomics of Infectious Diseases (CSGID). This project has been funded in whole or in part with U.S. Federal funds from the National Institute of Allergy and Infectious Diseases, National Institutes of Health, Department of Health and Human Services, under Contract Nos. HHSN272200700058C, HHSN272201200026C, and HHSN272201700060C

