## Supplementary matherial for Semper at al. for "Structural characterization of Nonstructural protein 1 from SARS-CoV-2"

**Table S1. Amino acid sequence similarity to SARS-CoV-2 Nsp1 of Nsp1 proteins from human pathogenic coronaviruses**

| Protein Name | Organism | % Identity to SARS-CoV-2 Nsp1 |
|---|---|---|
| MERS-CoV Nsp1 | Middle East respiratory syndrome-related coronavirus | 19 |
| HCoV-OC43 Nsp1 | Human coronavirus OC43 | 19 |
| HCoV-HKU1 Nsp1 | Human coronavirus HKU1 | 18 |
| HCoV-229E Nsp1 | Human coronavirus 229E | 9 |
| HCoV-NL63 Nsp1 | Human coronavirus NL63 | 6 |

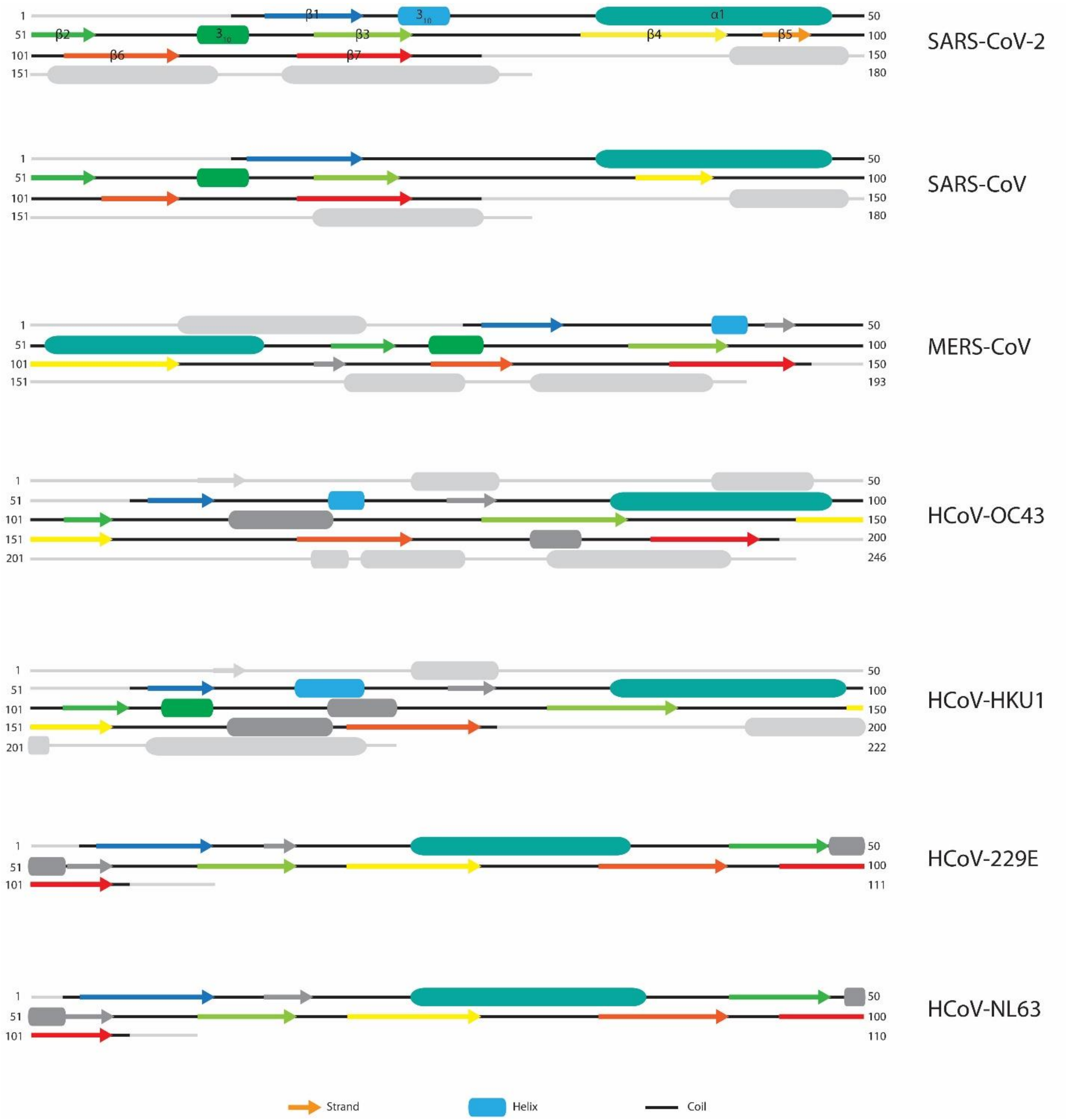


**Figure S1. Secondary structure analysis of Nsp1 from human-infecting Coronaviruses.** Within each sequence, the topological fingerprint containing elements that may facilitate formation of the capped β-barrel structure are coloured, while sequence outside of the fingerprint region is depicted in light grey. Key secondary structure elements observed in the SARS-CoV-2 Nsp1$_{13\text{-}127}$ crystal structure are labeled and are coloured accordingly in the ortholog sequences. Unique secondary structure features that lack an equivalent in the SARS-CoV-2 structure but that fall within the predicted β-barrel region are coloured as dark grey.
